# A geometric model of the visuomotor cortex as a sub-Riemannian assemblage of the visual and motor cortices

**DOI:** 10.64898/2026.08.06.743236

**Authors:** E. Baspinar, G. Citti, A. Sarti

**Affiliations:** MathNeuro Team, Inria Branch at the University of Montpellier, Montpellier, France; Department of Mathematics, University of Bologna, Bologna, Italy; Center for Social Analysis and Mathematics (CAMS), EHESS, CNRS, Paris, France

**Keywords:** Neurogeometry, visuomotor cortex, sub-Riemannian geometry, pinwheel

## Abstract

Classical neurogeometric models describe the primary visual cortex as a fibered structure in which retinal position and local orientation are coupled through the geometry of the roto-translation group. We extend this approach to the visuomotor cortex by modeling it as an assemblage of visual and motor cortical geometries. The model combines orientation-selective representations, analogous to those of the primary visual cortex, with movement-direction-selective representations, analogous to those of the primary motor cortex, in order to describe the mixed visual and motor selectivity observed in the visuomotor cortex. We introduce a coupled visuomotor structure in which visual orientation and motor direction coexist over a common spatial plane and interact through a relative-orientation constraint. Neural responses are modeled by orientation- and direction-dependent profile functions, and preference maps are obtained from vectorized population responses. Numerical simulations generate visual, motor, and mixed visuomotor response maps. A competition rule between visual and motor responses produces incidence ratios close to experimental observations in macaque visuomotor cortex. This framework provides a first neurogeometric approximation of visuomotor functional architecture and a mathematical setting for studying visually guided action.

## 1 Introduction

Humans and primates construct a perception of the surrounding environment throughout continuous interaction with the environment. This construction is often based on coordinated eye and arm movements. This suggests a cortical representation of such perception, a representation that involves visual, motor and visuomotor areas of the cortex.

Indeed, such a representation has been observed in certain cortical areas, in particular in the primary visual cortex (V1) [11], in part in the primary motor cortex (M1) [40, 29] and in the visuomotor cortex (V6A) [24, 25]. However, there is still very little understanding about how this cortical representation is constructed by the brain. Our goal is to provide a computational model for a better understanding of the way that the brain constructs this cortical representation in motor and visuomotor areas.

V1 is mainly responsible for identifying orientation alignments of the objects found in a 2D image. David H. Hubel and Torsten N. Wiesel showed experimentally that the orientation detection in V1 is done by specific neurons, which are organized in a columnar fashion [33, 34, 36, 32, 9, 10]. In each column, merely the neurons selective to the same orientation (preferred orientation, PO) are found.

There have been several experimental studies which suggest a similar columnar organization in M1 [40, 29]. M1 is mainly responsible for the execution of voluntary movements. M1 controls various features of motor behavior; in this work, we are interested in the direction of arm movement. While early studies demonstrated that individual neurons in M1 exhibit directional tuning [30], more recent investigations have shown that motor cortical activity reflects higher-level features of movement, including global movement features, movement primitives, and fragments of motor trajectories [31, 15]. However, in the present model, we consider only movement direction. This directional representation is based on specific M1 neurons, each of which is identified by its preferred (movement) direction (PD). These neurons are functionally organized in a columnar fashion in M1 with respect to their PDs, similar to the V1 neurons with respect to their POs [28, 50]. Nevertheless, there are still many ambiguities regarding the characterization of such functional organization in M1.

Although not as fine as in V1 and M1, V6A shows coarse retinotopy and strong modulation by arm movements [44]. This indicates a mixed selectivity rather than a purely sensory or purely motor selectivity based on columnar organization [12, 22]. Cortical representation of such mixed selectivity is still far from being understood. This calls for mathematical models to investigate the functional characteristics of V6A. It is particularly important that such models can be used to simulate experimental protocols, providing useful tools to refine existing approaches, and to design new experimental questions and methodologies.

Neurogeometric models have primarily focused on V1 [43, 16, 19, 7], with only recent extensions to M1 [38]. In [43], orientation selectivity of simple cells was modeled via a differential operator (one-form), forming the basis of a neurogeometric framework. This was further developed in [16], which used receptive profile filters that induce an anisotropic geometry on the rotation-translation group. The receptive profile filter was chosen as real or imaginary part of a Gabor function. This framework was later extended to incorporate scale [46, 47], as well as spatial frequency and phase selectivity [7], finally to temporal features of motion [3]. More recently, it has been applied to M1 to model PD selectivity and related activity patterns underlying arm movements [38].

In V1, the orientation preference map provides a 2D representation of the distribution of POs and pinwheels on the V1 surface (Fig. 1). The brain constructs this distribution by starting from the early postnatal period [53, 39, 37], and subsequently refines this distribution by using the activation patterns based on random retinal waves [51, 13]. This construction was modeled by superimposing random waves [41], and more recently, by using the neurogeometric framework in [16] and by following the experimental procedure which is employed to obtain orientation preference maps [20].

**Figure 1:**
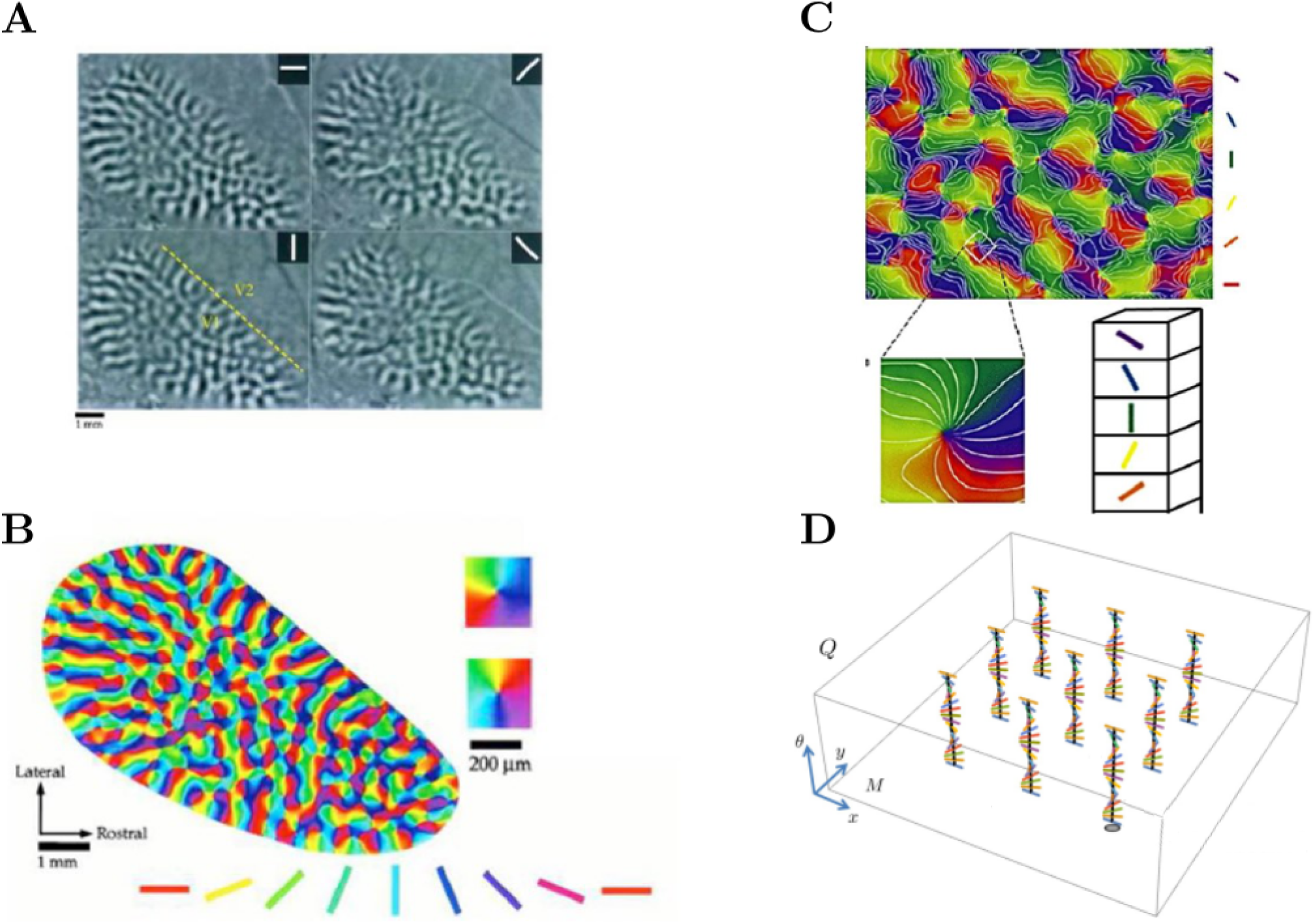
Orientation preference maps and pinwheels in V1. **A**. (Adapted from [11]) Responses of a tree shrew V1 to stimuli with four different orientations, of which each one is denoted at the top right of the corresponding recorded image. Black and white correspond to low and high response magnitudes, respectively. **B**. (Adapted from [11]) The orientation map obtained from the responses in A and represented in terms of a color map. Two pinwheels are highlighted on the right hand side. **C**. (Adapted from [42]) Illustration of an orientation map with isoorientation lines (white lines) and a highlighted pinwheel as the representation of a hypercolumn on the orientation map. **D**. Illustration of the geometric framework proposed for the V1 functional architecture. Each orientation fiber models a pinwheel composed of the simple cells sensitive to all possible orientations, which are denoted by the colored bars.

Similarly, in M1, direction preference map gives a 2D representation of the distribution of PDs on the M1 surface. In [40, 29], direction preference maps were obtained from monkey M1 via electrode recordings of the neurons involved in arm movement (Fig. 2A). These maps were analyzed and shown to provide a periodic pattern of repeated direction angles, similarly to the orientation preference maps of V1. Moreover, it was reported that the PDs between centers of repeating angles change gradually with respect to the distance as we move from one center towards another on the direction preference map. Based on these properties, a schematic lattice model of direction preference maps was proposed in [29] (Fig. 2B and 2C).

**Figure 2:**
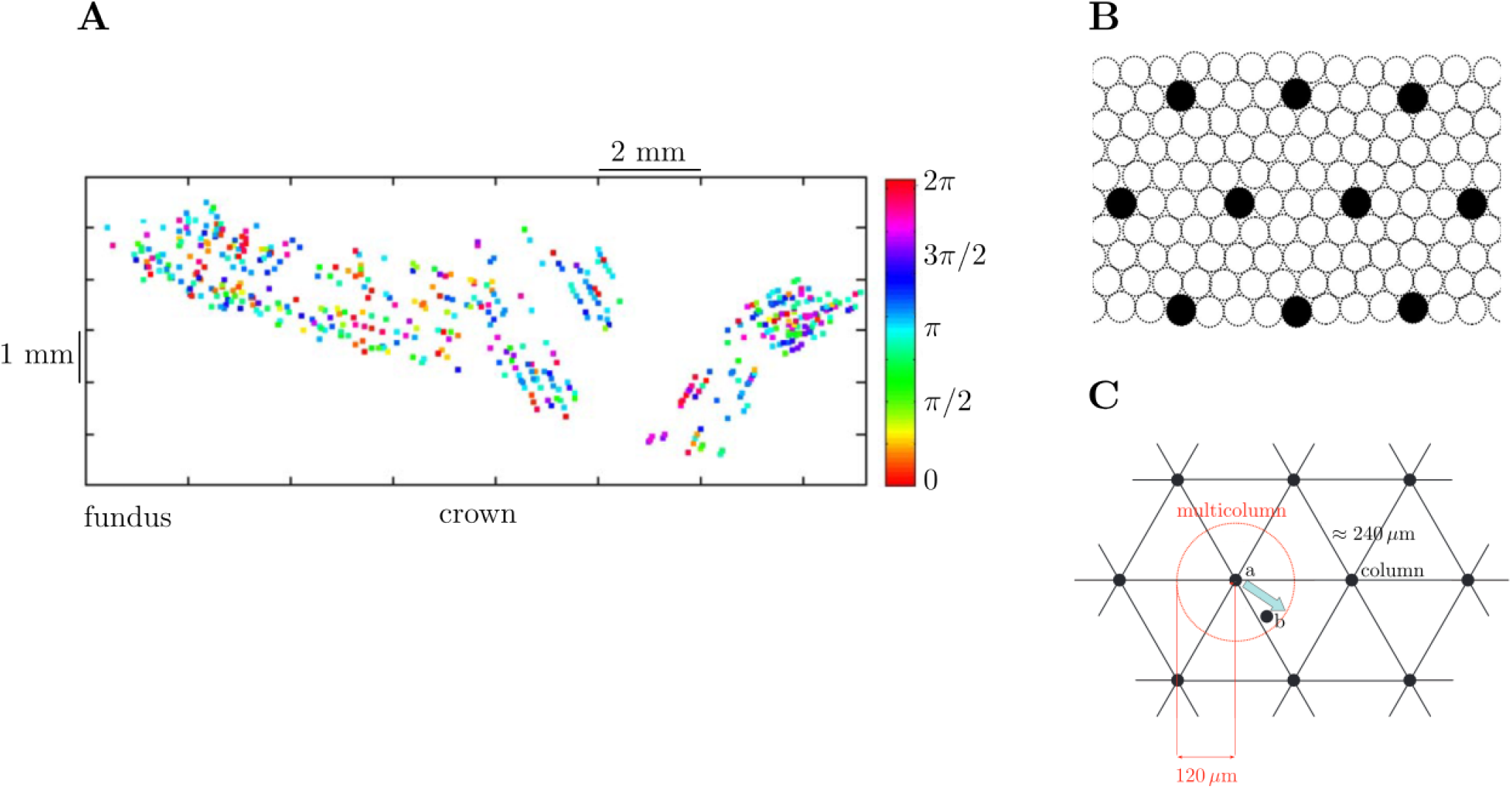
Direction preference map in M1. **A**. (Adapted from [40]) The direction map obtained from the neural responses recorded from monkey M1 during arm movements on a 2D plane. **B**. (Adapted from [29]) Schematic lattice model of the repeated, periodic mapping of PDs in M1. Each circle represents a column, in which merely the M1 neurons with the same PD are found. Black dots highlight the column locations highlighted in C. **C**. (Adapted from [29]) The same lattice model as in A. Column locations are highlighted. Red circle represents a multicolumn composed of the columns found in the disk region with a radius of 120 µm and centered around the column a. A multicolumn can be thought of as the M1 analogue of a pinwheel found in V1. The blue arrow shows the difference between the PD vectors of the neurons located at column a and column b. Spatial periodicity of the lattice is approximately 240 *µ*m.

In the present work, we extend the aforementioned previous computational studies of V1 and M1 to V6A. We propose a neurogeometric approach to study the cortical representation of selective nature of V6A. We start with a model for the construction of the direction preference map of M1 by extending the procedure which was proposed for the construction of V1 orientation preference map [5]. Then, we generate preference maps associated with the co-presence of both orientation and direction selectivity in V6A [24, 25]. We use a maximum selection procedure applied on the orientation preference and direction preference maps to obtain the preference maps of V6A.

The main challenge is that, differently from V1, M1 and V6A are not purely sensorial areas. Therefore, there is no concept of receptive profile in M1 and V6A. This is an obstacle to follow the approach in [16, 7] for the derivation of the neurogeometric frameworks regarding M1 and V6A. To tackle this, we invert the procedure given in [16], similarly to [43] but with a profile function. We first find an adequate one-form, and then design a profile function analogous to a receptive profile by using the one-form. A suitable candidate for such a profile function is the whole complex structure of Gabor functions, differently from the case in previous neurogeometric models of V1 [16, 7], in which only the real or imaginary component of the Gabor functions was used.

The Python code which produces the presented results is accessible in [6].

## 2 Neurogeometry

### 2.1 Neurogeometry of V1

V1 is a cortical area located in the occipital lobe of the brain. It is responsible for the first step processing of the visual input transmitted from the retina to the cortex. This first step processing focuses on the identification of orientation alignment of the objects in a 2D image.

Each neuron in V1 is responsive to a specific fragment in the 2D image, as well as to a specific orientation value at the location of this fragment. Its activity spreads across the whole network of V1 neurons. Through this interaction, the individual responses to local image fragments are integrated, resulting in a unified perception of the entire image.

David H. Hubel and Torsten N. Wiesel showed experimentally that orientation detection in a 2D image is achieved by the orientation sensitive V1 neurons, which are known as *simple cells* [33, 34]. They showed that the simple cells are organized based on *orientation columns* [34, 35]. *Each simple cell is* sensitive to a specific orientation value. In an orientation column, we find the simple cells which are sensitive to the same orientation value.

The orientation columns have a continuous arrangement across the cortex: POs of the neurons change gradually as we move across the cortex. This continuity is interrupted by the singularities on the cortical surface, which are known as *pinwheels* (Fig. 1B and 1C). On a pinwheel, in contrast to columns, we find the neurons that are sensitive to all orientations in the range of [0, *π*) [11]. That is, each orientation is represented equally as the others at the center of a pinwheel. The pinwheels are distributed periodically on the cortical surface, showing a crystalline-like arrangement [42, 17].

A pinwheel can be represented mathematically as an orientation fiber (Fig. 1D). According to this representation, the simple cells which respond to the same location in the 2D image, but to different orientations, are found in the same orientation fiber.

We represent an orientation fiber as located vertically at every point on V1 surface. Each point on V1 surface corresponds to a specific location on the 2D image plane. In this setting, V1 surface is represented as an identical plane to the 2D image plane. Identical mapping is assumed between image and retinal planes, as well as between retinal and V1 planes. This results in a one-to-one correspondence among the coordinates of these three planes.

It was proposed by Jean Petitot and Yannick Tondut, to model this configuration as a fiber bundle defined in a Heisenberg group [43]. This idea was further developed in [16], where the sub-Riemannian (sR) geometry of rotation-translation group was proposed as an adequate model of the V1 functional architecture. We consider this geometry as a smooth manifold, which we will denote by SE(2) ≃ ℝ ^2^ × *S*^1^.

Nearby neurons tend to be strongly connected, while distant ones are not. This reflects the anatomical connectivity based purely on spatial distance. However, neural interactions can also depend on response similarity: distant neurons tuned to the same stimulus may interact more strongly than nearby neurons tuned differently. This gives rise to functional connectivity, which depends on correlations in neural responses. A key example of such connectivity is long-range horizontal connectivity in V1, which is observed in ferrets, cats, and primates.

There is a high correlation between the neural responses of the simple cells which are sensitive to similar orientation values and to the fragments located at the same position in a given 2D image. The horizontal connectivity is strong between such highly correlated simple cells, whereas it is weak between the simple cells which are sensitive to different orientations, or to the fragments which are located at distant positions in the 2D image. This complexity requires a connectivity which integrates spatial and functional distances. This is very different from the anatomical connectivity, which is based only on spatial distance. The SE(2) geometry can generate the curves that closely approximate the long-range connectivity, and is therefore a suitable model for V1 functional organization.

In this geometric framework, we assume that the image (or external) plane and retinal plane are identical. We denote both by *M*, and the corresponding spatial variables by (*x, y*) ∈ *M*. We further assume that the retinal plane and cortical surface *M*_V_ of V1 are identical, i.e. *M* = *M*_V_ ⊂ ℝ^2^, thus (*x, y*) = (*x*_V_, *y*_V_) where (*x*_V_, *y*_V_) denote the spatial variables on *M*_V_.

Each simple cell in V1 responds to a particular spatial location (*x*_V_, *y*_V_) ∈ *M*_V_ and to a specific orientation *θ*_V_ ∈ [0, *π*) at that location. When an image fragment is present at (*x*_V_, *y*_V_) and oriented along *θ*_V_, the corresponding simple cell to (*x*_V_, *y*_V_, *θ*_V_) exhibits strong activation. In the absence of such a match, its activation is weak or even absent. This models the simple cell selectivity to both position and orientation. We refer to Section 3.1 for details.

The sR geometry of SE(2) is anisotropic: its metric induces a distance that combines both spatial and orientational components, as described in [16]. In this setting, we denote the columnar structure of V1 by *Q*_V_ ⊂ SE(2), where *Q*_V_ = *M*_V_ × [0, *π*)_V_. We denote each point in *Q*_V_ by (*x*_V_, *y*_V_, *θ*_V_), where *x*_V_, *y*_V_ are the spatial components and *θ*_V_ is the orientation component. Each point (*x*_V_, *y*_V_, *θ*_V_) ∈ *Q*_V_, denotes the location of the V1 simple cells which are sensitive to the specific position-orientation (*x*_V_, *y*_V_, *θ*_V_). We represent the neural activity at each such point as an *L*^2^-function *O*_V_ : *Q*_V_ → ℝ. In other words, *O*_V_(*x*_V_, *y*_V_, *θ*_V_) represents the activity of the neurons which are selective to the point (*x*_V_, *y*_V_) ∈ *M*_V_ and to the orientation *θ*_V_. We use the subscript *V* to highlight the differences between the frameworks of visual, motor and visuomotor areas in the cortex.

Simple cells that are sensitive to the same orientation *θ*_V_ are connected through long-range horizontal connections, even if they correspond to different spatial positions (*x*_V_, *y*_V_) ∈ *M*_V_ on the retinal plane. Consequently, interactions are particularly strong among simple cells sharing the same orientation preference and located within a sufficiently short distance of one another.

The long-range horizontal connectivity is approximated in SE(2) by the integral curves of the vector fields

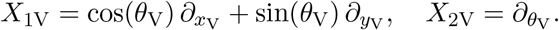

These vector fields are induced by the following one-form

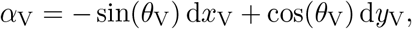

since

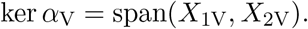

The vector fields *X*_1V_ and *X*_2V_ are called *horizontal* vector fields. The integral curves of these vector fields are called horizontal integral curves. We denote a horizontal integral curve with constant coefficients in *Q*_V_ by *γ*_V_ : [0, ∞) → *Q*_V_, which is given by

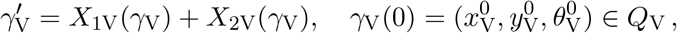

where *γ*_V_(*t*) is parametrized by time *t* and 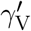 denotes the derivative of *γ*_V_ with respect to *t*. These horizontal integral curves provide the connectivity between the points in *Q*_V_.

As shown in [16], these horizontal integral curves approximate closely the psychophysical representations of long-range horizontal connectivity [23]. Neural activity in V1 tends to propagate primarily along these long-range connections [11]. Therefore, the sR geometry provides an appropriate framework for modeling activity propagation in V1.

### 2.2 Neurogeometry of M1

Directional tuning for movement is a basic functional property of the neural activity in M1. This directional tuning was first described for arm movements on a 2D plane [28]. Experimental studies showed that, the neurons which are tuned to a specific arm movement direction are found in the same column vertical to the cortical surface in M1 [29, 27]. On the contrary, neurons that are tuned to different movement directions are located in different columns. This suggests a columnar organization in M1, based on PDs (Fig. 2A and 2B).

This columnar organization is similar to the columnar configuration of orientation sensitive neurons found in V1 [34, 35]. In M1, we find a high correlation between the neurons with similar PDs and which are sufficiently close to each other [1, 8]. This is similar to the smooth change of POs across the V1 surface, as visualized in orientation preference maps (Fig. 1).

In V1, a hypercolumn has a pinwheel organization, which corresponds in an orientation map, to a singularity on which all preferred orientations are equally represented (Fig. 1). Despite the similarities to V1 columnar structure, in M1, such pinwheels in the form of singularities in the PD map have not yet been observed. Nevertheless, a disk-shaped organization, rather than pinwheels, was reported in M1 as potential structures that have the same function as pinwheels [40, 29]. This indicates that there might be a 2D representation of PDs in M1, similar to the orientation maps in V1, but which can be obtained at a different spatial scale. This is the motivation for us to extend the formalism proposed for V1 functional architecture [16] towards the columnar organization in M1 [38].

This approach focuses on the M1 area associated with arm movements on a 2D plane. It originates from the basic functional properties of M1 neurons tuned to movement directions. These properties involve tuning of hand position and movement direction [28].

In this approach, we denote the M1 cortical space by *Q*_M_ = *M*_M_ × [0, 2*π*)_M_ ⊂ SE(2), where *M*_M_ represents the base plane of M1. Similarly to the V1 case, we assume identical mapping between the external plane *M*, on which the arm movement takes place, and the surface, i.e. *M* = *M*_M_. We represent with the point (*x*_M_, *y*_M_, *θ*_M_) ∈ *Q*_M_, an M1 neuron that is tuned to a position (*x*_M_, *y*_M_) ∈ *M*_M_, and with PD equal to *θ*_M_. We represent the corresponding neural activity as an *L*^2^ -function *O*_M_ : *Q*_M_ → ℝ. Therefore, *O*_M_(*x*_M_, *y*_M_, *θ*_M_) represents the activity of the neurons which encode the arm movement at the point (*x*_M_, *y*_M_) ∈ *M*_M_ and in the direction of *θ*_M_.

In a perfect analogy with V1, the long-range horizontal connectivity in M1 is approximated by the integral curves of the vector fields

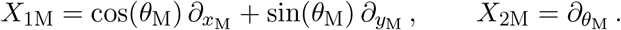

These vector fields are induced by the following one-form

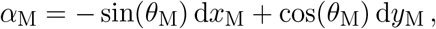

since by definition

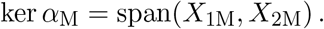

Similarly to V1, the integral curves of *X*_1M_ and *X*_2M_ are horizontal integral curves in *Q*_M_. We denote a horizontal curve with constant coefficients in *Q*_M_ by *γ*_M_ : [0, ∞) → *Q*_M_, which is described as

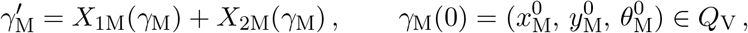

where *γ*_M_(*t*) is parametrized by time *t* and 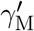 denotes the derivative of *γ*_M_ with respect to *t*. Within the model, these integral curves model the functional connectivity between the neurons found in *Q*_M_.

### 2.3 Neurogeometry of V6A

V6A contains: (i) visual neurons, responsive to object location and spatial features during eye fixation; (ii) motor neurons, active during reaching and arm movement execution; (iii) visuomotor neurons, which integrate both target position and movement parameters within the same cell. These neural classes are distributed in V6A in specific proportions. This distribution supports the interpretation of V6A as a sensorimotor interface rather than a strictly sensory or motor structure [24, 25, 12].

The neurogeometry of V6A can be considered as a synthesis of the geometric principles underlying V1 and M1. In V1, cortical organization reflects the geometry of visual space: retinotopy, orientation selectivity, and local feature integration define a structured manifold over retinal position and orientation. In M1, the geometry is expressed in terms of motor variables: position and directional tuning. Neurogeometry of V6A integrates these two frameworks, embedding perceptual and motor coordinates into a unified visuomotor geometry.

V6A can be described geometrically as a coupled visuomotor space, where visual and motor variables coexist within a unified structure rather than forming a simple fiber bundle. The visual component encodes both spatial location and local orientation, e.g. in gaze-centered or body-centered coordinates, while the motor component represents movement direction relative to the visual frame. Importantly, the interaction between these components depends on their relative orientation, preventing a strict separation into independent base and fiber variables. We denote this visuomotor geometry of V6A by *Q*_VM_ ≃ *Q*_V_ × *Q*_M_, emphasizing its product structure at the level of variables, while keeping in mind that the underlying geometry is intrinsically coupled. This is what is called *assemblage* [48, 49], *see Fig. 3*.

**Figure 3:**
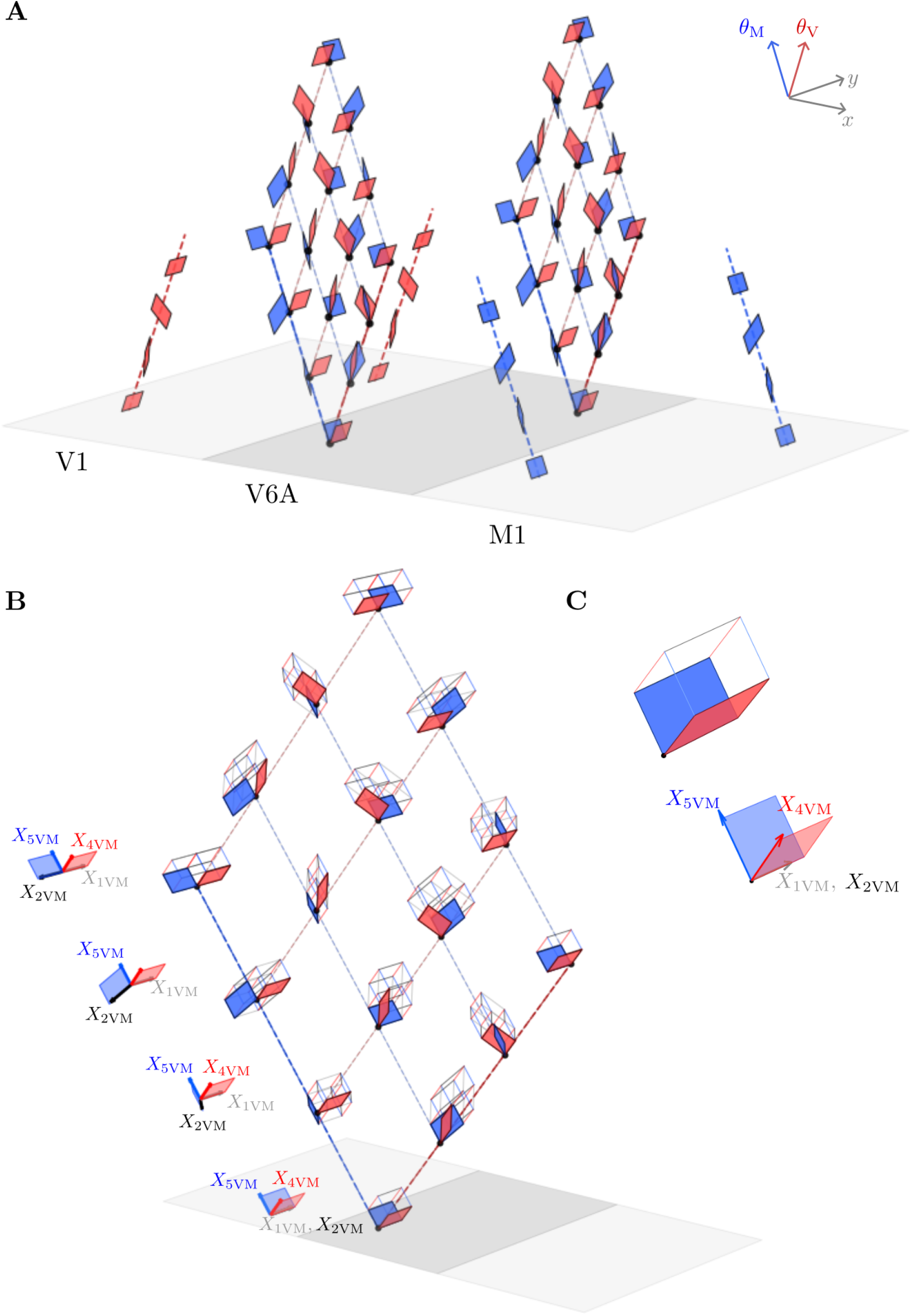
V6A as an assemblage of V1 and M1. Dashed red and blue lines denote visual and motor fibers, respectively; red and blue planes denote their tangent planes. Axes are shown in the upper-right corner. We assume (*x*_V_, *y*_V_) = (*x*_M_, *y*_M_) = (*x, y*). **A**. Orientation (V1, red) and direction (M1, blue) fibers, whose tangent planes rotate along *θ*_V_ and *θ*_M_, respectively. Center: 2D visuomotor (V6A) fiber obtained as an assemblage of orientation and direction fibers. **B**. Tangent planes on 2D V6A fibers form hypercubes. Left: vector fields spanning the four tangent planes associated with the first motor component of the 2D fiber. **C**. Degenerate case at the endpoints of the first visual component of the 2D V6A fiber. In this case, *X*_1V_ and *X*_1M_ coincide, causing the hypercube to collapse into a cube.

Unlike V1, where horizontal vector fields describe the propagation of orientation and position, and unlike M1, where vector fields represent movement direction in motor space, V6A requires horizontal vector fields acting simultaneously on both visual and motor components. Such horizontal vector fields generate sensorimotor trajectories which result from a coupling between visual and motor coordinates: a shift in perceived target position induces a corresponding displacement in motor configuration space. This coupling can be represented through a one-form that constrains the visual and motor coordinates:

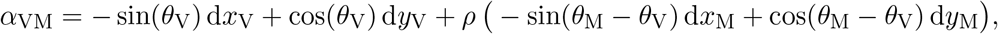

where the first and second components correspond to visual and motor parts, and where the motor part is multiplied with coefficient *ρ* > 0. Here *ρ* represents the coupling strength between the visual and motor components.

In this construction, the visual component *of α*_VM_ restricts the arm movement to directions aligned with *θ*_V_. The motor component encodes the arm movement relative to the orientation *θ*_V_. As a result of the dependence on *θ*_M_ − *θ*_V_, motor actions are encoded in eye-centered coordinates and expressed in a visual reference frame.

The one-form induces the following horizontal vector fields:

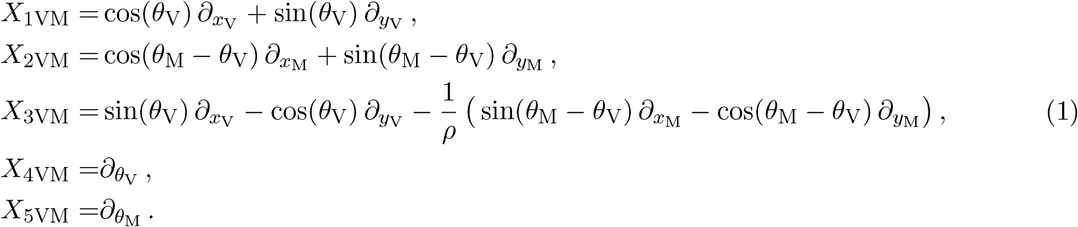

The corresponding geometric structure can be simplified by assuming

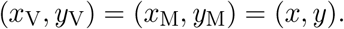

This is equivalent to assuming *M* = *M*_V_ = *M*_M_ in *Q*_VM_, ignoring the mappings between the external visual (or arm movement) plane and V6A base plane. Such mappings are complicated and out of scope of the present work; see [2, 14] for explanations about them. This reduced framework is illustrated in Fig. 3. Fig. 3A presents the geometric structures associated with the V1, V6A, and M1 fibers, together with the corresponding tangent planes as they rotate along the fibers.

The orientation fibers in V1 coincide with those of the classical neurogeometry of vision: the preferred orientation varies as one moves along the fiber in the *θ*_V_ direction. Similarly, the direction fibers in M1 can be regarded as the motor analogue of orientation fibers, with the preferred movement direction varying along the fiber in the *θ*_M_ direction.

In V6A, the fibers are two-dimensional and arise through an assemblage of orientation and direction fibers. This construction is fundamentally different from simply juxtaposing the orientation and direction fibers, as illustrated in Fig. 3B. Instead, the tangent planes associated with the 2D visuomotor fibers are organized into four-dimensional hypercubes, whose edges are aligned with the vector fields generating the V6A geometry, i.e. with the four vector fields given in (1).

Pairs of tangent planes on a 2D visuomotor fiber encode the gaze direction and the hand-movement direction associated with a given spatial location, corresponding to the base point of the fiber in the (*x, y*) plane. As shown in Fig. 3C, a degenerate configuration occurs at the endpoints of a 2D visuomotor fiber. At these points, the vector fields *X*_1VM_ and *X*_2VM_ coincide, causing the associated four-dimensional hypercube to collapse into a three-dimensional cube. This configuration is mathematically degenerate because the distribution loses one independent direction. From a neurophysiological perspective, this degeneration corresponds to the case in which gaze and hand movement are aligned and share the same direction (or orientation), so that the eyes follow the trajectory of the hand.

A horizontal integral curve with constant coefficients in this setting is denoted by *γ*_VM_ : [0, ∞) → *Q*_VM_. Within the model, it approximates the functional connectivity between the neurons in *Q*_VM_. It is described by

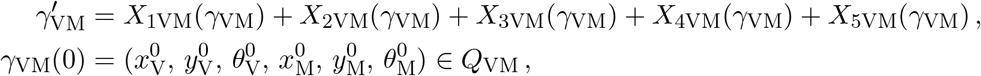

where *γ*_VM_(*t*) is parametrized by time *t* and 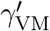 denotes the derivative of *γ*_VM_ with respect to time.

## 3 Modeling neural responses

Neural response profile refers to the impulse response function of a neuron. In the case of visual neurons, this response is induced by a perceptual (visual) input. It allows the neuron to extract visual features from the visual input, such as orientation. In the case of motor neurons, the response is induced by a cortical stimulation. It allows the neuron to generate a motor action from the received cortical stimulation.

### 3.1 Response to visual input

Gabor functions have been considered as suitable mathematical representations of the impulse response functions of the simple cells. This is justified by the fact that they provide the optimal compromise between space and orientation localization [21, 4].

We use the compact notation 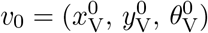 to denote a point *v*_0_ ∈ *Q*_V_. We will use the following coordinates to simplify the notation:

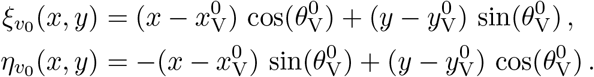

Then, we denote by 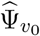 the real part of a Gabor function centered at 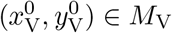 and rotated by 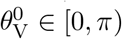

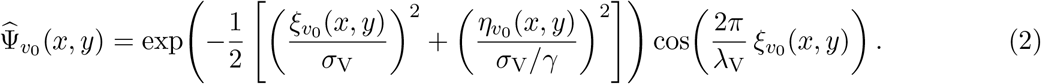

Here *γ* is a constant in (0, 1]. It is the ratio between horizontal and vertical components of the Gabor function. It becomes more sharply selective to a specific orientation as it decreases from 1. The constant *σ*_V_ is strictly positive and it determines the scale of the Gaussian localization. Large scale values model the neurons sensitive to the features of larger fragments whereas it is the opposite for small scale values. Finally, *λ*_V_ *>* 0 is a constant that determines the wavelength. Small wavelength models the neurons selective to high spatial frequency whereas it is the opposite for high wavelength.

We model the visual neural response, i.e. receptive profile of the orientation-sensitive visual neurons, as the following normalized Gabor component:

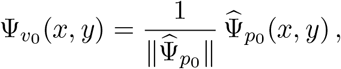

where ∥ · ∥ denotes the standard Euclidean norm.

The response of the simple cell sensitive to the position 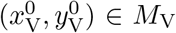 and to the orientation 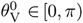, is found via the following convolution with the cortical stimulation denoted by an *L*^2^-function *I* : *M*_V_ → ℝ:

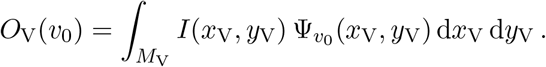

This representation provides a scalar measure of a single neuron activity. We use the vectorized sum of the simple cell responses [54, 52] so as to provide a representation of the population activity induced by the simple cells sensitive to the position 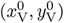, but sensitive not necessarily to the same orientation.

We vectorize the scalar responses *O*_V_ to represent which orientation each response corresponds to:

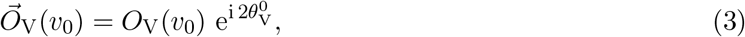

where 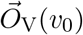 denotes the vectorized version of *O*_V_(*v*_0_). Double-angle is needed in the vectorizing exponential since the exponential is the covering map *θ* ⟼ e^i2*θ*^, which maps from the circle *S*^1^ to the real projective line *RP* by gluing the antipodal points on the circle. This mapping avoids that opposite directions cancel each other since an orientation does not have any direction.

We use the following vector sum to find the population response of the neurons at 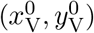

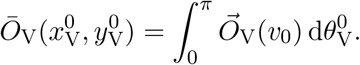

Finally, we find the orientation which the neurons at 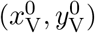 are sensitive to via

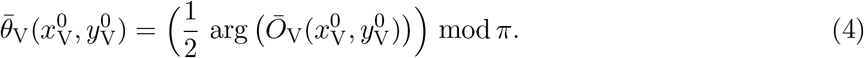

This provides the orientation angles represented in orientation preference map; see Fig. 4A. Dividing by 2 is needed to compensate the double-angle used in 3.

**Figure 4:**
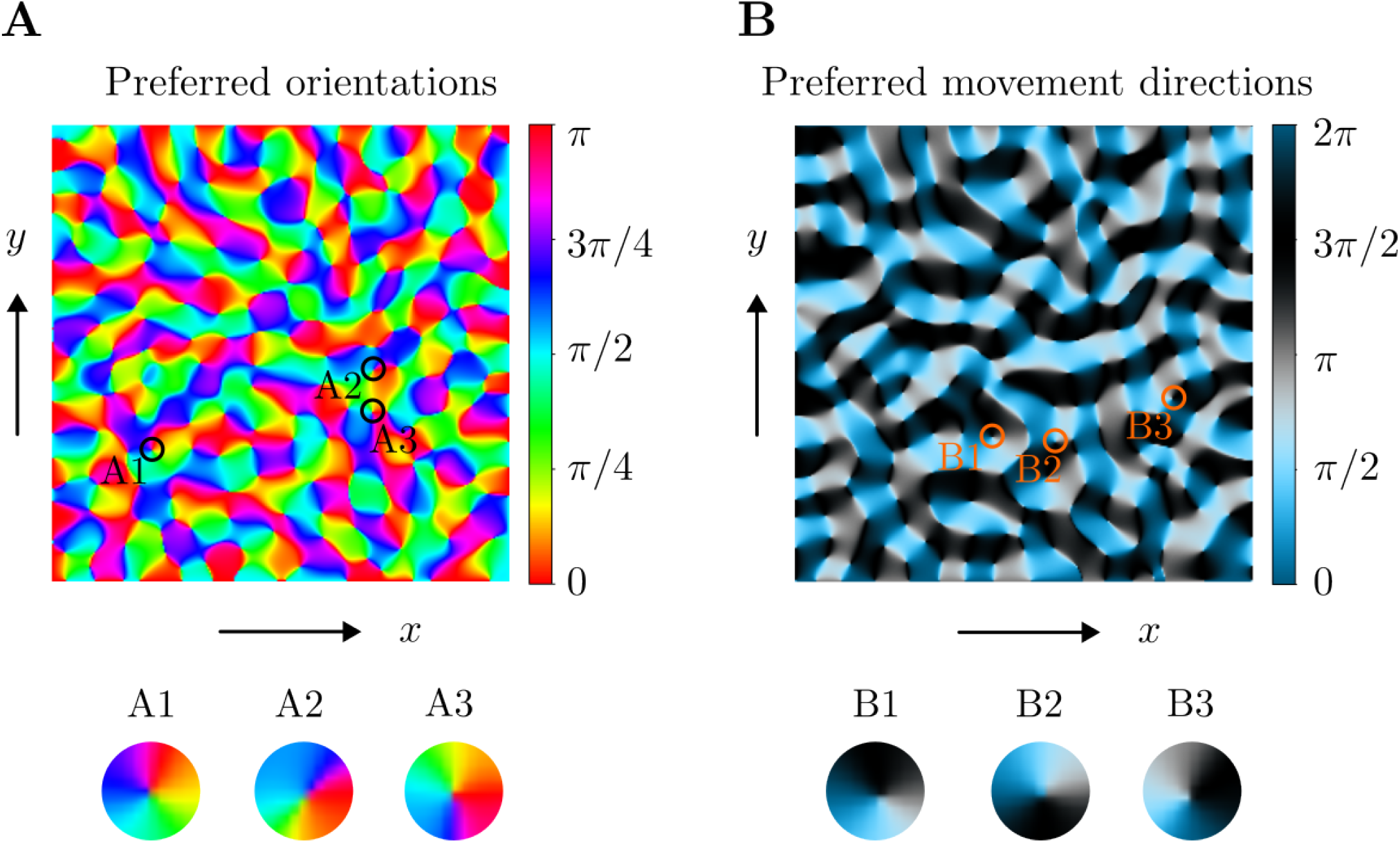
Preference maps with pinwheels. Maps of preferred orientations (A) and preferred movement directions (B) are obtained from the noise image. The vertical bars provide the colormaps for the orientations and movement directions. Below each preference, three highlighted pinwheels (A1, A2, A3) and multicolumns (B1, B2, B3) are provided as zoomed in.

### 3.2 Response to motor input

We use the notation 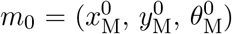 to denote a point *m*_0_ ∈ *Q*_M_. We follow the same idea as in the case of visual input. The only difference is that we use the odd component 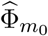 of Gabor function to model the response profile of motor neurons at *m*_0_:

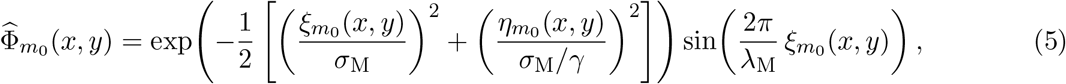

and the response profile is given by

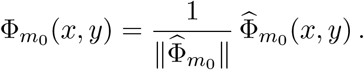

Stimulated activity of the motor neuron which controls the movement at the point 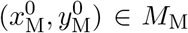 and with preferred direction 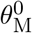, is found via

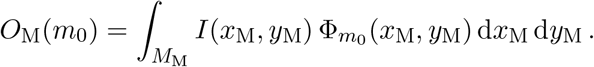

Here *I* : *M*_M_ → ℝ is an *L*^2^-function that models the cortical stimulation, analogously to the visual stimulation in the case of visual neural response.

Following the same line of thought used in the case of visual input, we vectorize the scalar responses *O*_M_ via

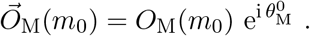

An important difference compared to (3) is that the exponential does not have double-angle because now the direction is taken into account. Differently from orientation, which is *π* − periodic, direction is a 2*π* − periodic quantity. Therefore, no mapping to projective line is required.

In arm movements, a large population of motor neurons is involved during movements in different directions [30, 26]. This suggests that the same neural population is likely to be involved in generating movements in various directions. If a neuron responded only to its preferred direction, activation alone of such cells with this preferred direction would be enough to generate a movement in that direction. However, since their tuning is broad, individual neurons respond to multiple directions, so movement direction is encoded collectively by the activity of a population rather than individual neurons alone. This also provides robust responses, similar to each other in the cases where the same input stimulates the neurons.

We model such collective encoding of movement direction 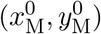 as follows:

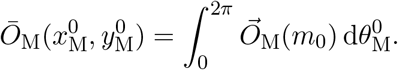

We find the movement direction to which the neurons at 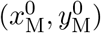 are sensitive to, via

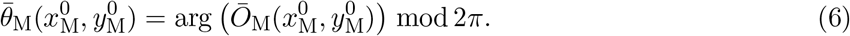

This provides the direction angles represented in direction preference map; see Fig. 4B.

### 3.3 Response to visuomotor input

We model the neural response to a visuomotor input in V6A as the maximum between the visual and motor responses evoked by that input. This maximum-selection rule assumes that, at any given location in V6A, the neural population (visual or motor) producing the stronger response to the visuomotor stimulus, determines the selectivity of that site – whether visual (perceived orientation) or motor (movement direction). This assumption is motivated by the fact that both neuron types are found on the V6A surface, in a mixed way. Therefore, they are likely to receive the same input as cortical stimulation of V6A. However, it is possible that they do not respond in the same way to all stimuli [24, 25].

From [14], we deduce that there are two types of change of variables:

- a change of variables from (*x*_V_, *y*_V_) ∈ *M*_V_ to external coordinates (*x, y*) ∈ *M*,
- a change of variables from (*x*_M_, *y*_M_) ∈ *M*_M_ to external coordinates (*x, y*) ∈ *M*.

We assume for simplicity that the visuomotor input is mapped on external coordinates (*x, y*) ∈ *M*. It is an *L*^2^-function denoted by *I* : *M* → ℝ. For each (*x, y*) of the external plane *M*, we consider the coordinates (*x*_V_(*x, y*), *y*_V_(*x, y*)) ∈ *M*_V_, which are the visual coordinates corresponding to the same point (*x, y*) on the external plane. If these coordinates are not unique, we consider the following set:

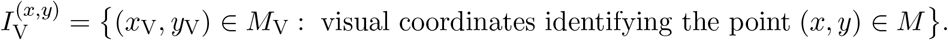

We can consider the motor coordinates (*x*_M_, *y*_M_) ∈ *M*_M_ in the same way for each point (*x, y*) ∈ *M*. If these coordinates are not unique, we can consider the following:

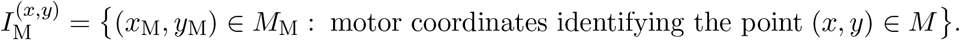

We will assume that the visual and motor coordinates are unique for each point (*x, y*) on the external plane *M*.

We model the neural response *O*_VM_ : *M* → ℝ to the visuomotor input as follows:

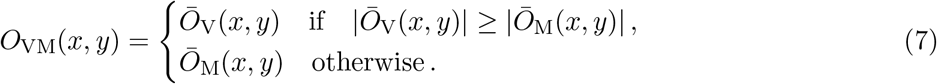

This allows us to trace which origin the neural response comes from. In this way, we can construct the distribution map of visual and motor neurons in V6A. We refer to Fig. 5A and 5B for examples of experimentally observed and numerically generated distribution maps of V6A, respectively.

**Figure 5:**
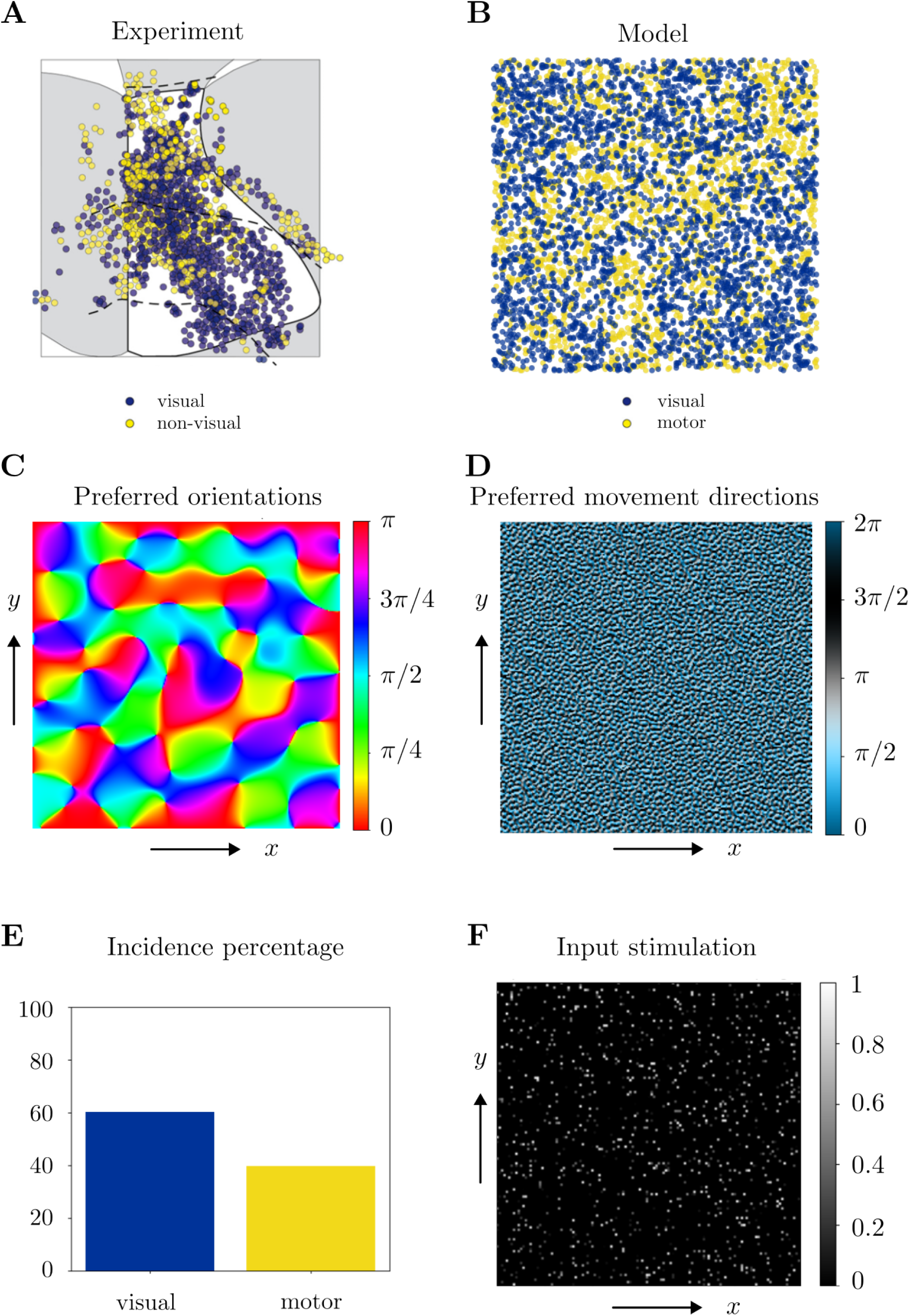
Distribution of visual and motor responses, preference maps and incidence percentages. **A**. (Adapted from [25]). Experimentally observed distribution of visual (blue) and non-visual (yellow) neurons in V6A. **B**. The distribution obtained via the model simulations by using (7). **C**. and **D**. Preference maps obtained via (4) and (6). **E**. Incidence percentages obtained from B. **F**. Noise image used as the input stimulation for visual and motor neurons in the simulations of Fig. 5B, 5C and 5D.

## 4 Results

### 4.1 Model generates realistic preference maps

We model the visuomotor input as a 256 × 256 sparse random noise image *I* : *M* → [0, 1). To generate the stimulation pattern, we generate a 256 × 256 probability array, in which each entry corresponds to a probability value sampled from a uniform distribution over [0, 1). Each entry in this array is associated to a pixel of the visuomotor input. Finally, we fix a probability threshold, which we denote by *p*. Each pixel in the visuomotor input is then stimulated if its associated probability entry is smaller than *p*: each entry is compared to *p*. If it is smaller than *p*, the associated pixel is marked as true (stimulated); otherwise, it is marked as false (non-stimulated). The pixel intensity value of each pixel which is marked as true is independently drawn from a uniform distribution over [0, 1). We assign directly the intensity value 0 to the pixels which are marked false.

We provide in Fig. 4, the orientation and movement direction preference maps which we obtain by using the Gabor responses to visual and motor inputs, respectively. We fix *p* to 1, i.e. all pixels are stimulated. In the Gabor components given by (2) and (5), we choose *σ*_V_ = *σ*_M_ = 16, *λ*_V_ = *λ*_M_ = 32 and *γ* = 0.8. We calculate preferred orientations and movement directions by using (4) and (6), respectively. We represent the preferred orientation and direction angles on the 2D plane in terms of the corresponding colormaps provided in Fig. 4A and 4B. In Fig. 4A and 4B, we provide the zoomed in images of three highlighted pinwheels and three multicolumns, respectively.

### 4.2 Model provides insight: visuo-motor competition in V6A

Experimental studies show that the visual and non-visual neurons in V6A are distributed with a certain ratio, and apparently in a chaotic manner, without any specific order. In [24], 61% of these neurons are reported to be visual, and 39% to be non-visual. A similar incidence percentage of these neurons is provided in [25], as well as their distribution map in V6A; see Fig. 5A.

In the experiments performed in [24, 25], visual and non-visual neurons are identified in terms of to which stimulation type they respond to. Visual, somatosensory and motor inputs were tested. The neurons which provided strong response to visual input were considered as visual neurons. The neurons which did not provide any noticeable response to the visual input, but a strong response to somatosensory and/or motor inputs, were classified as non-visual.

Our simulation results provide approximate results of these experimental observations; see Fig. 5. These results suggest that the construction of V6A functional organization is based on a competition between visual and motor activity. It is important to note that we do not make any difference between somatosensory and motor neurons in the model. We consider all motor neurons as non-visual, ignoring the neurons that are purely somatosensory.

We provide in Fig. 5B, the distribution of visual and motor responses generated by our model. The response types in the model-generated distribution are found by using (7). We use the 256 × 256 test image given in Fig. 5F. We set *p* = 0.059 for the probability of pixel stimulation. The following Gabor parameters are used: *σ*_V_ = 32, *σ*_M_ = 2, *λ*_V_ = 2 *σ*_V_, *λ*_M_ = 2 *σ*_M_, *γ* = 0.8. The results were obtained by using (7) and by randomly downsampling the points: each point computed via (7) is kept with a probability of 0.12, the rest is ignored and not shown in the distribution map. The corresponding incidence percentages of visual and motor responses are given in Fig. 5E. The incidence percentages (60% visual, 40% motor) were obtained after the downsampling.

In Fig. 5C and 5D, we provide the preference maps which were used to obtain the visual and motor responses given in Fig. 5B. The orientation and movement direction preference maps were obtained from the same test image via (4) and (6), respectively. On the contrary to the distribution map in Fig. 5B, there is no downsampling applied to the preference maps given in Fig. 5C and 5D.

### 4.3 Model produces a large variety of cortical maps

We provide in Fig. 6 the incidence percentages of the visual and motor responses with respect to changing *p* values used in generating the input stimulation. In this result, we use 256 × 256 images which were generated with varied *p* values between 0 and 1. The Gabor parameters are the same as those used in Fig. 5B. We compute the incidence percentage for each *p* value by following the same procedure as in Fig. 5E.

**Figure 6:**
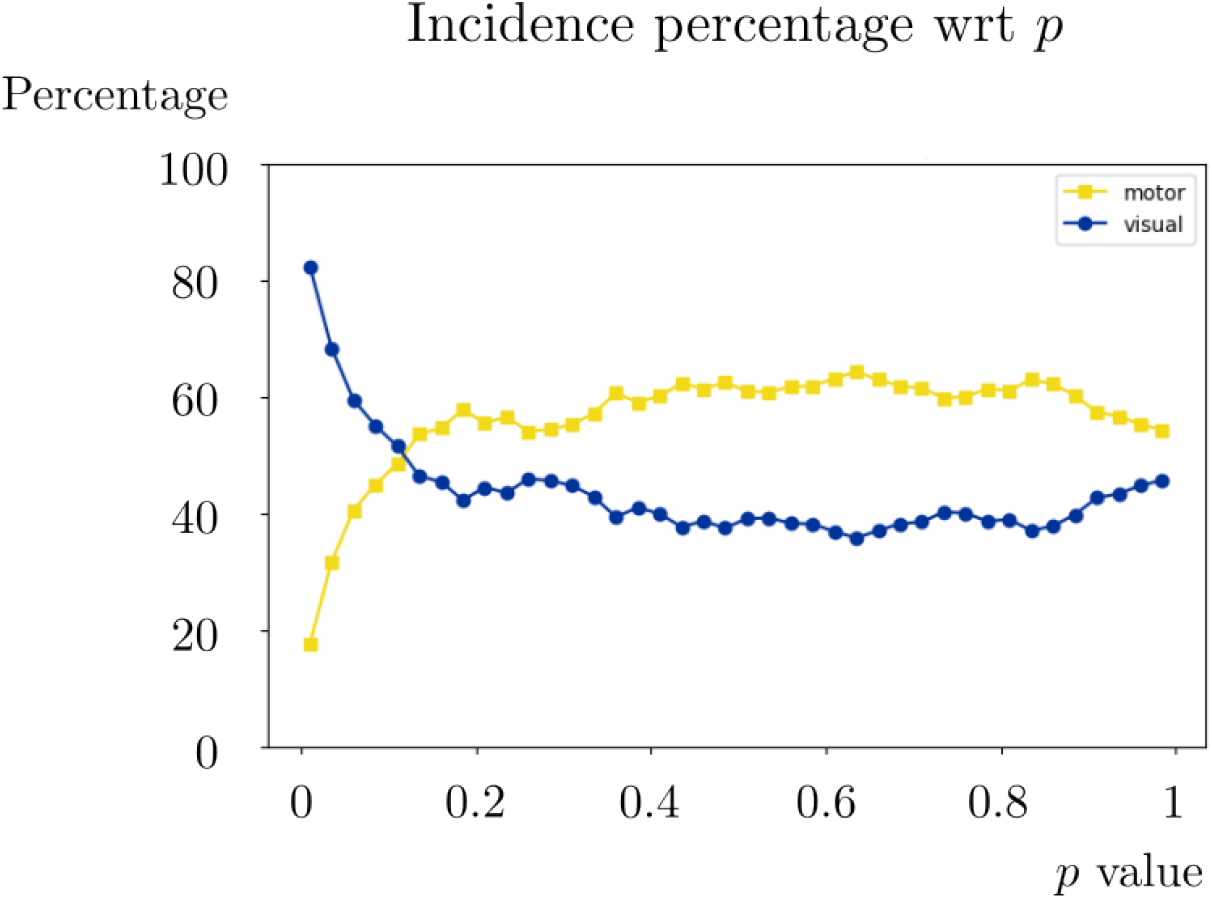
Incidence percentages of visual and motor responses with respect to probability *p*.

The model can generate a variety of incidence percentages, making it adequate to study individual differences among different subjects in the experiments. Nevertheless, it is important to choose a sufficiently small value for probability *p* since beyond *p* value ≈0.125, motor responses have a higher incidence percentage than the visual responses. Such a high incidence of motor responses is not consistent with the experimental observations given in [24, 25].

In Fig. 7, we provide the preference maps which are obtained at different scales: *σ*_V_ = *σ*_M_ = 8, 16, 32. We use the same 256 × 256 input image which was used in Fig. 4, with *p* = 1. Gabor parameters are *λ*_V_ = 2 *σ*_V_ and *λ*_M_ = 2 *σ*_V_ for each scale value. Finally, *γ* = 0.8.

**Figure 7:**
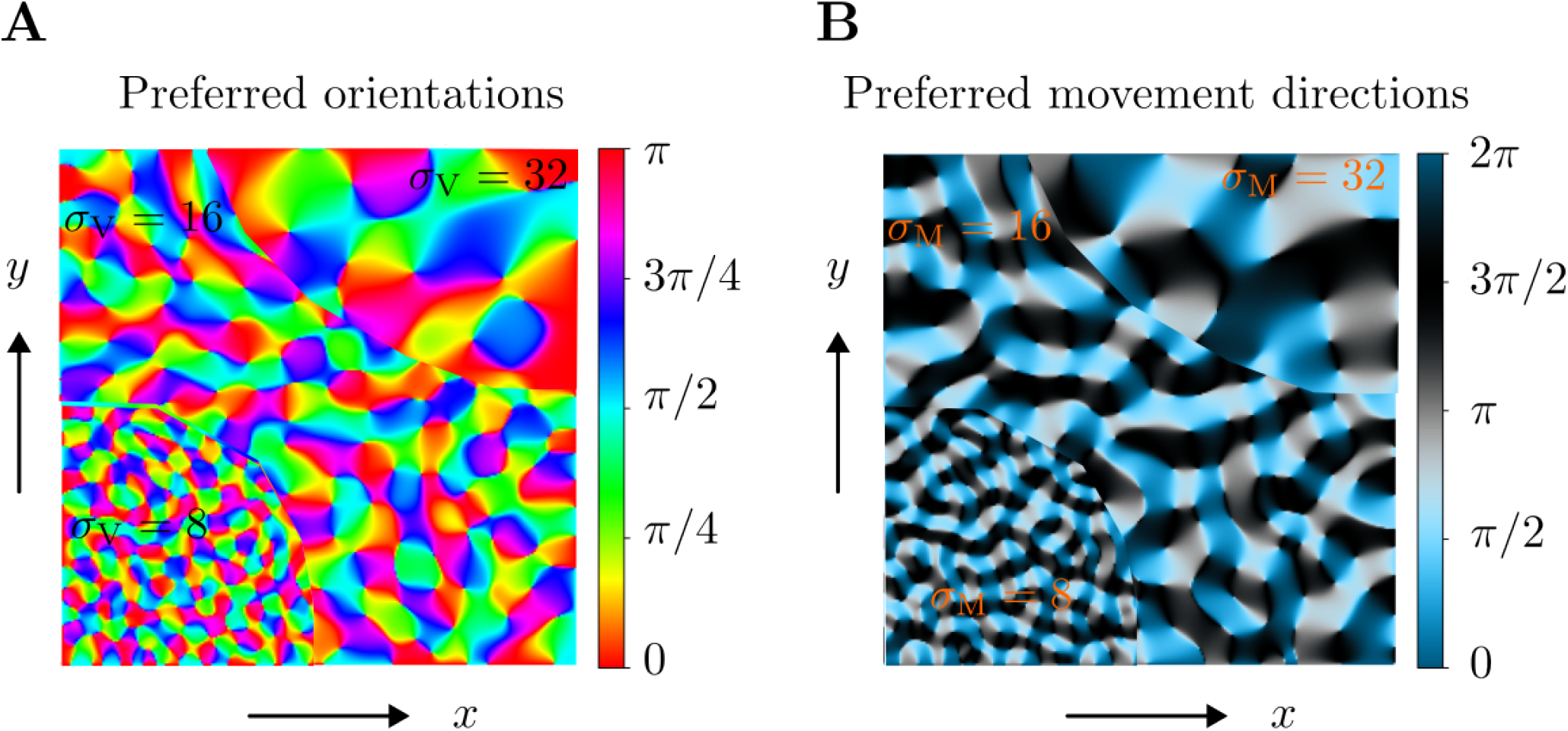
Preference plots obtained by using Gabor filters with scales *σ*_V_ = *σ*_M_ = 8, 16, 32.

In the distribution maps of neural responses to visuomotor input, an important parameter which determines the incidence percentage is the scale values *σ*_V_, *σ*_M_ and their ratio *σ*_V_*/σ*_M_. In Fig. 8, we provide the corresponding distribution maps and the incidence percentages for varying scale ratios. We observe that the incidence percentage of visual responses decreases and that of motor responses increases, as *σ*_V_*/σ*_M_ ratio decreases.

**Figure 8:**
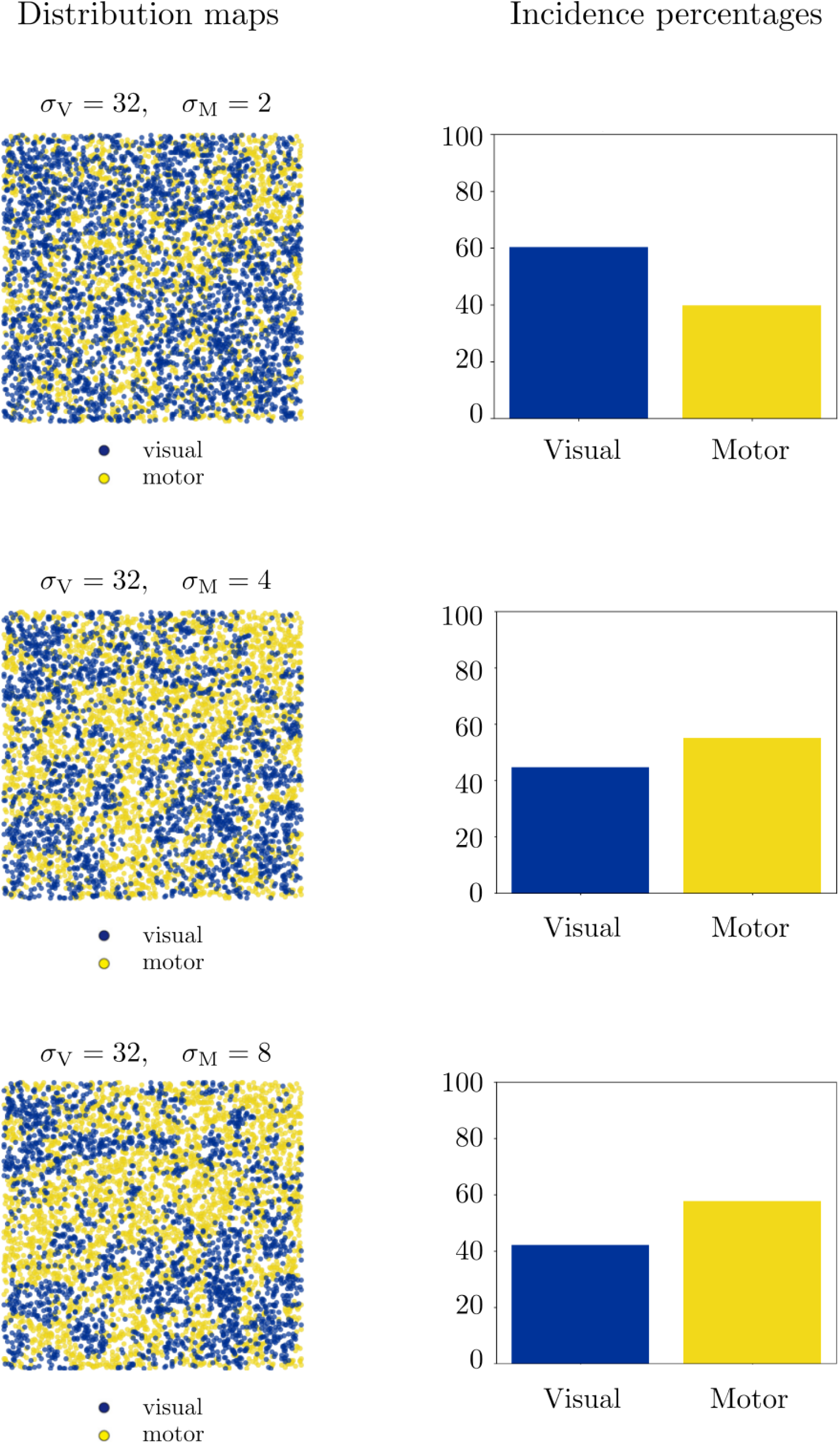
Distribution maps and incidence percentages of visual and motor responses. Responses are obtained from the same visuomotor input used in Fig. 5F. We vary the scale values of *σ*_M_ from 2 to 8 by keeping *σ*_V_ fixed to 32.

These results show that our model is potentially flexible to be fitted to a variety of experimental results obtained from different subjects, which may exhibit individual differences in their cortical structures.

## 5 Discussion

In this work, we proposed a neurogeometric model of the visuomotor cortex V6A by extending the classical neurogeometry of vision to a sensorimotor setting. The proposed framework combines the geometric principles previously developed for V1 and M1 into a unified visuomotor structure. More precisely, we introduced V6A as an assemblage of visual and motor geometries, in which orientation-selective and movement-direction-selective representations coexist within a common mathematical framework. This construction provides a first geometric approximation of the functional architecture of a cortical area characterized by mixed visual and motor selectivity.

The model builds directly upon the neurogeometric theory originally proposed for V1 [43, 16, 42]. In these models, cortical organization emerges from the geometry of the roto-translation group SE(2), where position and orientation are represented as coupled variables. Such a framework successfully explains several properties of V1, including orientation selectivity, pinwheel organization, long-range horizontal connectivity, and contour integration [43, 41, 16].

More recently, similar geometric principles were extended to M1, where preferred movement direction replaces preferred orientation as the principal tuning variable [38]. Our work represents a further step in this progression by addressing cortical areas that cannot be characterized as purely sensory or purely motor. Instead, V6A contains neurons with visual, motor, and genuinely visuomotor properties, requiring a geometric structure that simultaneously incorporates both visual and motor representations.

A central novelty of the present model is the introduction of a coupled visuomotor geometry. Rather than treating visual and motor variables independently, we defined a one-form whose structure explicitly depends on the relative orientation between visual and motor variables. This leads to horizontal vector fields that simultaneously govern the evolution of visual and motor coordinates and generate sensorimotor trajectories in the combined space. The resulting geometry is therefore not a trivial product of the V1 and M1 geometries, but rather a coupled structure in which visual and motor representations influence one another. Such coupling reflects the experimental observation that V6A participates in transformations between visual information and arm movement parameters during visually guided actions [24, 25, 12].

A second conceptual contribution is the introduction of 2D visuomotor fibers obtained through the assemblage of orientation fibers and movement-direction fibers. In the classical neurogeometry of vision, a fiber represents the set of neurons associated with a single retinal location but selective to different orientations [43, 16]. Similarly, in the M1 framework, fibers represent neurons associated with a movement location but selective to different movement directions [38]. In contrast, the fibers proposed here for V6A are intrinsically 2D and simultaneously encode visual orientation and movement direction. Their tangent spaces are naturally organized into 4D hypercubes generated by the visual and motor horizontal vector fields. This geometric interpretation provides a novel representation of visuomotor selectivity and suggests a possible mathematical substrate for the integration of sensory and motor information within a common cortical architecture.

The computational results further support the relevance of the proposed framework. First, the model generates orientation and movement-direction preference maps exhibiting organizational principles analogous to those reported experimentally in V1 and M1, respectively. In particular, the simulations reproduce pinwheel-like organization for orientation selectivity and multicolumn-like organization for movement-direction selectivity. These results indicate that similar geometric mechanisms may underlie the emergence of preference maps in both sensory and motor cortices.

Second, the model reproduces experimentally observed distributions of visual and non-visual neurons in V6A. Using a simple maximum-selection rule between visual and motor responses, the simulations generate mixed distributions whose incidence percentages closely approximate those reported experimentally [24, 25]. The emergence of these distributions suggests that the heterogeneous organization of V6A may arise from a competition between visual and motor representations. Although simplified, this interpretation offers a possible mechanistic explanation for the mixed selectivity observed experimentally and provides a quantitative framework for investigating how sensory and motor representations interact within the same cortical area.

An additional strength of the model is its flexibility. By varying stimulation statistics and geometric scale parameters, the framework generates a broad spectrum of cortical maps and incidence ratios. This flexibility is particularly relevant in light of the substantial inter-individual variability observed in cortical organization. The possibility of fitting different experimental datasets through changes in a small number of geometric parameters suggests that neurogeometry may provide a useful low-dimensional description of complex cortical architectures.

A limitation of the present work should nevertheless be emphasized. Our geometric setting assumes an identical spatial base plane for visual and motor representations, i.e. (*x*_V_, *y*_V_) = (*x*_M_, *y*_M_) = (*x, y*). This simplification allows us to focus on the geometric coupling between orientation and movement direction while avoiding the complexities associated with visuomotor coordinate transformations. However, it is very likely that visual and motor representations are expressed in different reference frames and are related through nontrivial transformations [2, 14]. Therefore, our model should be regarded as a first-order approximation of the underlying sensorimotor geometry rather than a complete description of V6A organization.

A future research direction will therefore be the introduction of distinct visual, motor, and visuomotor spatial coordinates. Instead of using a common spatial base plane, the geometry could be generalized to incorporate separate coordinate systems. This would transform the current simplified assemblage into a richer structure which can be used to better understand how visual information is progressively encoded into motor commands.

Another future direction is the derivation of neural activity propagation equations directly from the proposed horizontal connectivity structure. This will require an extension of the equations which were proposed in the V1 cortical geometry [45] to the V6A geometry. Such developments would provide a model of visuomotor connectivity, which is inevitable to extend the activity propagation in V1 [16, 18]. This could enable the simulation of dynamic visuomotor processes, including target selection, reach planning, and eye-hand coordination.

Overall, the proposed model extends neurogeometry beyond the classical domain of visual perception and toward the study of sensorimotor integration. By combining the geometric principles of V1 and M1 into a coupled visuomotor framework, it provides a mathematically grounded description of V6A functional organization and offers a new perspective on how visual and motor representations may coexist within a common cortical geometry. Finally, the model suggests that neurogeometry may provide a unified language for understanding functional architectures across sensory, motor, and sensorimotor cortical systems.

## Appendix: Discretization

We now describe the discretization used in the numerical simulations. The same notation is used for visual and motor responses. We write

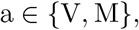

where a = V denotes correspondence to V1 and a = M denotes correspondence to M1. The angular domain has period

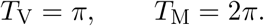

For a fixed number *N*_*θ*_ of angular samples, angular step size is 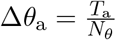 . The discretized angles are

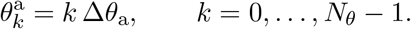

In the simulations, we use *N*_*θ*_ = 64.

The input stimulation is represented by a discrete image

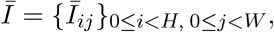

with *H* = *W* = 256 in the simulations. The sparse random stimuli are generated as

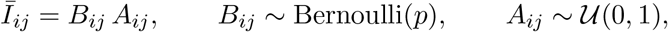

independently for all pixels (*i, j*), with *U*(0, 1) representing the uniform distribution over [0, 1).

For each modality a, each angle 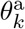, and each discrete offset (*m, n*) in the finite filter support, we define the rotated coordinates

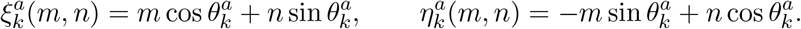

Finite filter support is defined by

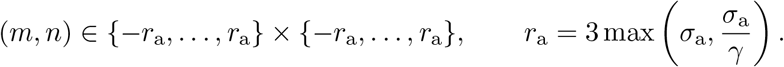

The discrete Gabor profile is

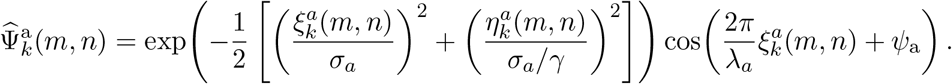

The phases are chosen as

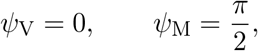

so that the visual response is obtained from the even Gabor component, while the motor response is obtained from the odd Gabor component. Each discrete kernel is normalized by its Euclidean norm,

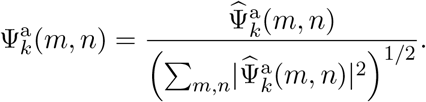

This corresponds to the normalization used in the Python implementation.

The discrete neural response is then computed by filtering the image with every angular kernel:

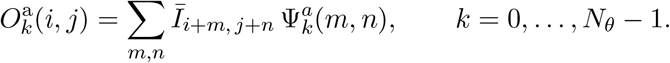

Thus the numerical lifting of the image is the array

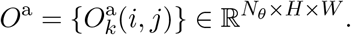

This is the discrete version of the continuous responses *O*_V_ and *O*_M_. In the code this is implemented by applying `cv2.filter2D` to the image for all angular kernels.

The population vector at each pixel is obtained by summing the angular responses with complex angular weights. We use

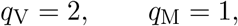

where the factor 2 for V accounts for the fact that visual orientation is *π*-periodic, while motor direction is 2 *π*-periodic. The discrete population response is

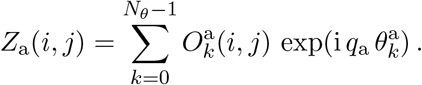

The preferred angle and response magnitude are then

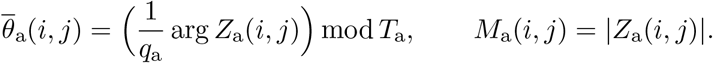

For a = V, this gives the preferred visual orientation map; for a = M, it gives the preferred movement direction map.

The visuomotor selection is made pixelwise by comparing the vectorized response magnitudes:

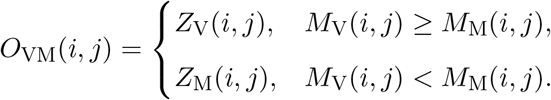

Equivalently, the response type assigned to the pixel (*i, j*) is visual when

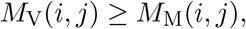

and motor otherwise.

